## Supplementary Figure 1 for "Thermal Cycling-Hyperthermia Attenuates Rotenone-Induced Cell Injury in SH-SY5Y Cells through Heat-Activated Mechanisms"

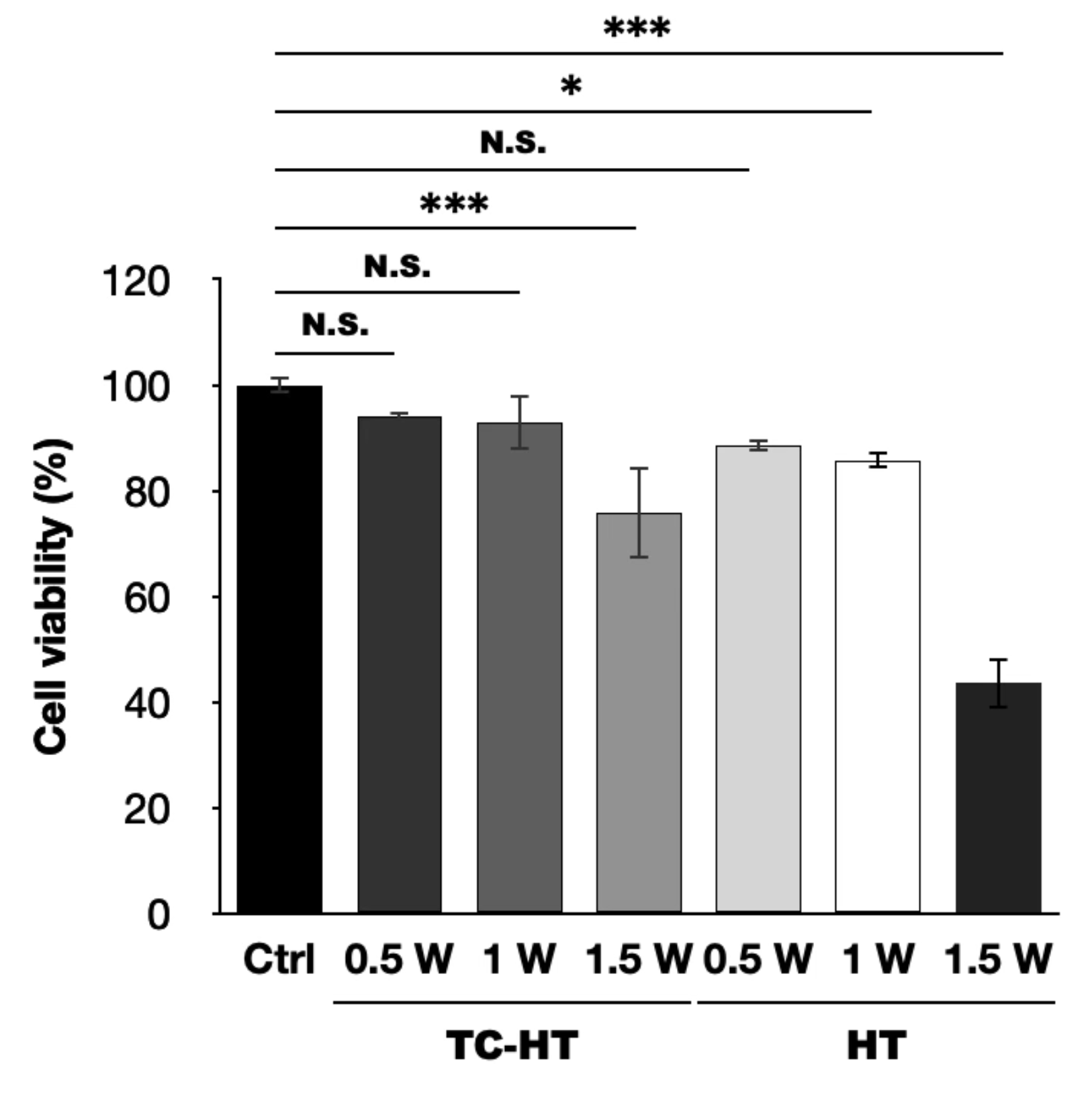

**Supplementary Figure 1.** Neurotoxicity of TC-HT and HT at various intensities used alone on SH-SY5Y cells. CCK-8 assay was conducted to determine the viabilities of SH-SY5Y cells following TC-HT and HT treatments alone with different US intensities. The viabilities were measured 24 h after TC-HT and HT treatment. Data were presented as the mean ± standard deviation in triplicate. Significance levels between indicated groups are denoted as *P < 0.05 and ***P < 0.001, while non-significant differences are indicated as N.S.
